# Genomic insights into the human gut commensal *Megasphaera elsdenii*: Relatedness and metabolic potential compared to animal isolates

**DOI:** 10.64898/2026.07.30.741745

**Authors:** Dina Sabirova, Mikhail Rayko, Veronika Vinichenko, Anton Shikov, Radick Altinbaev, Kirill Antonets, Milyausha Yunusbaeva, Bayazit Yunusbayev

## Abstract

*Megasphaera elsdenii* is best known as a prominent lactate consumer within the rumen microbial community in livestock, and its metabolic properties are relatively well studied. In humans, it can be isolated from healthy donors’ feces and, more often, from patients’ feces with diverse inflammatory conditions. Genetic diversity of this species is poorly understood, and it is currently unclear whether human and animal gut isolates are genetically related and perform the same metabolic function. In this study, we compared 86 *M. elsdenii* genomes from human feces (as a proxy for the human gut) to those of animal gut isolates. Phylogenetic analysis revealed that human and animal gut lineages intermingle within a single, genetically homogeneous branch, lacking any host-specific clustering. Human gut lineages shared most of their genes and biochemical pathways with those of swine and cattle gut isolates, despite differences in their digestive tracts. Neither unsupervised nor supervised approaches identified any notable differences in encoded pathways between different host-specific gut lineages. Genome-scale metabolic modeling suggests that human and animal gut lineages likely share identical carbon and energy source requirements. Moreover, the requirements for lactate and acetate were conserved across all studied samples, regardless of the host. Finally, we found no virulence genes, and lactate utilization remains a plausible explanation for *M. elsdenii* accumulation in the host intestine.

**IMPORTANCE:** *Megasphaera elsdenii* is considered a commensal in the human gut and animal rumen. However, *M. elsdenii* tends to be more abundant in patients’ feces with diverse inflammatory conditions. As we know little about the strain diversity and biology of human gut lineages, comparisons with better-studied animal isolates can be informative. In this study, we compared human gut *M. elsdenii* genomes to those from better-studied isolates from ruminant and non-ruminant animal hosts. Human gut samples associated with patients and healthy donors were genetically very similar to gut isolates from animals and may have shared a common origin. We found that human and animal gut lineages have a similar genomic makeup and metabolic potential, and that neither group harbors virulence genes. We hypothesize that *M. elsdenii* is a benign commensal that grows in response to lactate accumulation in the inflamed gut.

## INTRODUCTION

*Megasphaera elsdenii* is best known as a prominent lactate-utilizer in the rumen and a fatty acid producer, and it has recently gained attention in human gut microbiome studies. This gram-negative anaerobic bacterium from the *Veillonellaceae* family has a long history of research (over 70 years) in animal microbiology [1]. From the very beginning, studies documented its ability to ferment lactate and produce C2–C6 volatile fatty acids, such as propionate, butyrate, and valerate, as well as carbon dioxide and hydrogen. Starting with the earliest works, *M. elsdenii* was studied for its ability to consume lactate and stabilize rumen acidity [2]. Rumen acidity can be caused when animals are fed rapidly fermentable substrates, and lactate production exceeds utilization by the rumen microbial community. Rumen acidity and ruminitis (inflamed rumen mucosa) have an adverse impact on the host, leading to laminitis, liver abscesses [3], and mortality [4]. While *M. elsdenii* can be isolated in diseased animals, its capacity to metabolize different forms of lactate has a beneficial effect on the host, proving it to be a promising probiotic [5, 6, 7]. Later meta-analyses have supported the beneficial effects of this bacterium on the host [8, 9]. While ruminal strains are better studied and likely beneficial, lineages associated with the human gut are less well studied.

### *Megasphaera elsdenii* form human lower gastrointestinal tract

In human microbiome studies, microbial communities inferred from fecal samples often serve as a proxy for the microbial community of the lower gastrointestinal tract, denoted as colon microbiota or, more often, gut community. Thus, *M. elsdenii* can be isolated from the fecal material of healthy individuals [10, 11, 12], but it occurs at a higher abundance in the feces of patients having diverse inflammatory conditions, such as acute HIV [13], CADASIL arteriopathy [14], colorectal cancer [15], and psoriasis [16, 17]. Its presence in healthy individuals, but higher abundance in patients, can have the same explanation as its distribution among healthy and diseased livestock. Namely, in ruminant animals, *M. elsdenii* can be present in a healthy rumen, but increases in abundance in response to lactate accumulation and therefore correlates with lactic acidosis and rumen dysbiosis [18]. In support of this explanation, recent experiments showed that *M. elsdenii* from human feces utilize lactate and co-occur with lactate-producers, such as *Lactobacillus* and *Streptococcus thermophilus* [12].

For a long time, *M. elsdenii* has been considered a human gut commensal [10]. However, a recent study revisited this traditional view by exploring immunostimulatory properties of fecal isolates from patients with colorectal cancer (CRC). *M. elsdenii* derived from CRC patients triggered dendritic cell (DC) maturation and activation via LPS-induced TLR4 signaling when gavaged to antibiotic-treated mice and Specific-Pathogen-Free mice [15]. While this study described activation-associated changes in murine immune cells, no evidence was presented for immunogenicity, i.e., the adaptive immune response to *M. elsdenii*, which would require both signal 1 (microbial antigen) and signal 2 (conserved microbial patterns, such as LPS) [19]. In other words, there were no antibodies specific to *M. elsdenii* or clonal T cell expansion with TCR specificity to *M. elsdenii*. Overall, the study findings boil down to evidence that *M. elsdenii* from CRC patients have lipopolysaccharides (LPS) that can stimulate antigen-presenting cells. As such, the study findings are consistent with similar evidence for ruminal *M. elsdenii* isolates, i.e., typical immunostimulatory capacity of microbial-derived LPS [20]. We note that relatively high immunostimulatory capacity was registered for vaginal phylotypes [21], but vaginal *Megasphaera* phylotypes are currently considered a distinct species and are tightly associated with vaginosis [22].

Taken together, there are fragments of data suggesting that *M. elsdenii* and related species can occur in health and disease, cell wall components (LPS) can be immunostimulatory, and some gut lineages resemble ruminal isolates in metabolic capacity. At the same time, it is unclear whether health and disease associations of this species can be explained by strain heterogeneity and whether different strains have detectable differences in genetic makeup and function. Finally, we lack a comprehensive understanding of the evolutionary relationships and genetic diversity among gut-associated lineages from different organisms [23]. To address these questions, we employed comparative genomics to analyze *M. elsdenii* samples associated with the human gut, together with better-studied gut isolates from ruminant and non-ruminant animals. We compiled *M. elsdenii* sequences representing healthy donors and psoriasis patients and published assembled genomes derived from animal gut isolates, as well as human vaginal phylotypes for context. We assembled *M. elsdenii* genomes derived from human fecal metagenomes and analyzed their phylogenetic relationships with isolates from the cattle rumen, swine intestine, and human vagina. By reconstructing metabolic pathways from genomic data, we predicted metabolic potential shared by all the studied *M. elsdenii* lineages and then sought differences between different hosts. We did not find notable differences in metabolic profiles between gut-associated lineages from humans, swine, and cattle. Instead, our findings show that the potential to utilize lactate, acetate, and simple sugars is encoded in all studied lineages and represents a unifying carbon source for this species. Finally, our analyzes suggest that human gut lineages lack virulence genes and are likely commensals that grow in response to lactate accumulation in the intestine.

## MATERIALS AND METHODS

### *M. elsdenii* from psoriasis patients’ fecal material

A case-control study focused on the gut microbiome of psoriasis patients found that *M. elsdenii* was one of the four human gut bacteria (*Megasphaera elsdenii*, *Catenibacterium mitsuokai*, *Eubacterium sp. CAG:180*, and *Rothia mucilaginosa*) associated with psoriasis [17]. This study also showed that *M. elsdenii* and three other top disease-associated species correlated with increased levels of fecal calprotectin, a biomarker of intestinal inflammation. This study hypothesized that low-grade inflammation in patients’ colon leads to lactate accumulation and *Megasphaera elsdenii* growth. For our current study, we examined metagenomic sequences from 47 healthy adult donors and 53 psoriatic patients. For genome assembly, we considered metagenomic sequence data from 13 donors with elevated relative abundance of *M. elsdenii* and sequencing reads (> 50000) sufficient for genome assembly [17]. Metagenomic sequences were quality-filtered and decontaminated as described in the original study [17]. Briefly, FastQC v0.11.9 [24] was used for sequence quality control, and Trimmomatic v0.33 [25] was used to clip Illumina adapters and perform quality trimming. Tandem Repeats Finder v4.09 was used for tandem repeat trimming [26]. Finally, KneadData v0.10.0 from the bioBakery toolkit was applied to decontaminate reads originating from the human genome, transcriptome, and microbial RNA [27].

### Genome assembly strategies

To assemble *M. elsdenii* genomes, we used metaSPAdes v4.0.0 [28], which is designed to handle metagenomic data. To obtain metagenomically assembled genomes (MAG), we used two approaches. The binning-based approach to obtain MAGs was implemented using MetaBAT 2 v2.12.1 [29] by setting the following parameters (minContig 2500, minCV 1.0, minCVSum 1.0, maxP 95%, minS 60, maxEdges 200). The obtained bin qualities were assessed based on completeness, contamination, and ribosomal marker presence using CheckM v1.1.6 [30]. A reference-based approach to obtain MAGs was implemented using Minimap2 v2.24 [31]. The QUAST package v5.2.0 [32] was used to assess the quality of the genome assembly based on N50, L50, total assembly length, number of contigs, longest sequence, GC-content, assembling errors, and unique and repeated region analysis. Assembled contigs were united into scaffolds using RagTag v2.1.0 [33].

### Reference genome choice for read mapping approach

To choose a reference genome for our analyses, we considered three fully assembled genomes of *M. elsdenii* downloaded from the NCBI database. Two reference genomes (strains NCIMB702410 and ATCC25940) were isolated from cattle, and one (strain 14-14) from a swine (Table S1). To choose a reference genome, we assessed overall genome relatedness among three available reference genomes using average nucleotide identity (ANI) [34]. High OrthoANIu values (> 97.72) indicated that the compared genomes were highly similar, and all three could be used for read mapping (Table S2). Nevertheless, the GCA_003006415.1 genome (strain NCIMB702410 from cattle) was preferred because it was assembled using SPAdes. A reference genome assembled with a similar assembler yields better read mapping accuracy. Finally, by taking GCA_003006415.1 as a reference, we mapped other genomes and visually inspected the spatial distribution of matching and non-matching regions using BRIG v1.0.0 (BLAST Ring Image Generator) [35]. Mapped genomes (red and green circles) had numerous visible gaps, indicating structural variants (Fig. S1).

### Metagenomically assembled genomes using a reference-based approach

Genome assembly approaches can vary in performance depending on the characteristics of the sequencing dataset. To select the most accurate approach, we compared the two alternative methods for assembling genomes using a single metagenomic sequence (from human donor D68, denoted as MAG68 in Table S3). Our comparative analysis showed that the reference-based approach performed better. For example, in sample 68, we retrieved 116 (93.5%) BUSCO genes using the binning approach and 122 (98.4%) using the reference-based approach. Hence, to obtain MAGs, we used the reference-based approach on our full dataset of 13 metagenomes. MAG quality and completeness were evaluated using BUSCO v5.5.0 (Benchmarking Universal Single-Copy Orthologs) [36]. For downstream analyzes, we selected 11 MAGs with BUSCO scores > 70% (arbitrary threshold; Fig. S2). We validated these MAGs using CheckM; they demonstrated completeness of 63-100% and contamination of < 6% (except MAG63), which corresponds to medium-quality draft according to the MIMAG reporting standards, i.e., completion ≥ 50%, contamination < 10% [37]. For reference, high-quality draft should have ≥ 90% completion, and < 5% contamination. After obtaining MAGs, their contigs were assembled into longer scaffolds using RagTag [33].

### Annotating genes and pathways

After obtaining 11 MAGs with longer scaffolds, we annotated coding sequences (CDS) and inferred gene functions using Prokka v1.14.5 [38]. The annotated genes were also classified into KEGG pathways using the FACoP.v2 tool (Functional Annotation and Classification of Proteins of Prokaryotes) in FUNAGE-Pro [39].

### Constructing pan genome for *Megasphaera* species

To infer the core versus the accessory genome of the analyzed set of *Megasphaera* species, we constructed a pan genome using Roary v3.11.2 [40] based on Prokka-annotated data for MAGs. Next, we used the Phandango v1.3.1 tool to visualize the pan genome [41], resulting in a heatmap showing filled and empty bins for genes that are either present or absent in the accessory part of the pan genome. To further analyze host groups in the pangenome, we used MicrobiomeAnalyst v2.0 [42].

### Phylogeny based on orthologous genes

To infer phylogenetic relationships between *M. elsdenii* lineages and related species, we first obtained a supermatrix of orthologous genes for all the analyzed genomes using BUSCO [43]. A maximum likelihood (ML) phylogenetic tree was then inferred based on this supermatrix using IQ-TREE v2.3.1 [44]. The ML tree was then visualized using ITOL v6 (Interactive Tree Of Life) [45].

### Analysis of KEGG pathways and CAZymes

*M. elsdenii* isolates from different hosts can have different metabolic adaptations, and KEGG pathways can have nonrandom distribution among different lineages. To understand how metabolic (KEGG) pathways are structured across hosts, we applied unsupervised methods, such as distance-based clustering and principal component analysis (PCA). Specifically, KEGG pathway abundances were converted to distances (Jensen–Shannon divergence) and grouped using Ward clustering, as implemented in the ‘Clustering analysis’ module of MicrobiomeAnalyst v2.0 (Fig. 3B) [42]. KEGG pathways were also analyzed using PCA within the same module to find major axes of pathway variation in the sample set (Fig. 4B). Finally, we manually defined host-associated classes and applied a supervised approach to seek differences. To determine KEGG pathways that explain differences between predefined classes (e.g., animal vs. human gut), we used linear discriminant analysis with effect size estimation (LEfSe) [46]. We next characterized CAZymes (Carbohydrate-Active enZYmes) in *M. elsdenii* genomes by searching for homologs from the CAZy database (accessed November 15, 2025) [47] using the MMseqs2 v14.7e284 [48]. A 70% identity and mutual coverage threshold was used to call hits in the database. We counted the total number of genes encoding CAZymes in each screened genome and then tested for differences in the median number of CAZymes harbored by gut lineages from humans, swine, and cattle. Median differences were tested using the Wilcoxon paired test, followed by the Benjamini-Hochberg (BH) procedure for multiple-comparison adjustment. The total number of CAZymes in each group (humans, swine, and cattle) was then classified into different CAZyme families. Proportions of the 10 most frequent families were analyzed using ANOVA (Analysis of Variance) followed by Tukey’s Post-Hoc Test with BH correction. Finally, we used MOB-suite software tools to detect plasmids in the 25 draft genomes that were used for metabolic predictions (KEGG, CAZymes) [49]. We could not detect (reconstruct) plasmid sequences in the 25 draft genomes (Table S3).

### Genome-scale metabolic modeling with gapseq

For the combined set of assembled genomes and MAGs (N=25), we predicted encoded metabolic pathways, reactions, and transporters using gapseq [50]. Predicted pathways, reactions, and transporters were used to build a draft model and predict a growth medium, assuming an anaerobic condition (removing oxygen from the final medium).

### Search for virulence factor genes and structurally similar proteins

To search for virulence factors in the *M. elsdenii* genomes, we used the MMseqs2 tool version v14.7e284 [48]. Known virulence factors were based on the VFDB 2022 (Virulence Factor Data Base) [51], accessed May 28, 2024. For each query, the best hit from the database was selected based on the e-value, a statistical parameter that quantifies the expected number of hits by chance when searching a database of a particular size. We retained the best hits that covered more than 70% of the query sequence, using a 70% identity threshold. In theory, known virulence factors from distant species can be missed due to sequence divergence. In the meantime, proteins from distant species can retain similar structure and function despite sequence divergence [52]. We therefore tried to identify virulence factors using a protein-language model implemented in the VirulentHunter tool, which can capture structural (and possibly functional) similarity. VirulentHunter employs a pre-trained ESM2 protein-language model [53] that was fine-tuned on proteins associated with virulence factors [54]. In VirulentHunter, the ESM2 protein language model was fine-tuned on virulence factors from three databases: VFDB 2022 [51], Victors [55], and BV-BRC [56]. When we first applied VirulentHunter to our *M. elsdenii* dataset, we were puzzled by the sheer number of “predicted virulence factors”, reaching approximately 200 proteins per genome. To interpret VirulentHunter predictions for *M. elsdenii*, we performed similar predictions for a phylogenetically related gut commensal (*Dialister hominis*) and true pathogen (*Clostridium botulinum*).

## RESULTS

### Human gut-derived *M. elsdenii* show little divergence from animal gut isolates

To assess genetic diversity and perform comparative analyzes, we retrieved publicly available *Megasphaera elsdenii* genomes from NCBI (Table S3), representing human and animal gut (cattle and swine) isolates (accessed March 18, 2024). Publicly available genomes were combined with 11 metagenomically assembled genomes (MAGs), generated in this study, representing *M. elsdenii* from psoriasis patients experiencing low-grade inflammation and one healthy donor. MAGs, unlike genomes from isolates, represent a consensus across a microbial population present in the analyzed fecal sample [30]. We, therefore, use “*M. elsdenii* lineage or population” when we refer to MAGs and when we refer to MAGs in combination with published genomes derived from isolates. For phylogenetic analyzes, we also included genomes of outgroup species, and other *Megasphaera* species, as well as two *Megasphaera* phylotypes from the human vagina [57], which were recently classified as distinct *Megasphaera* species [22]. We quality-filtered (BUSCO score > 70%, small genome size) our initial collection (N=98) and retained 86 genomes (Table S3) to explore genetic similarities between *M. elsdenii* and related species using ANI. While some isolates are denoted as separate species, genetic similarities, as indicated by ANI values, were greater than 95% and, in some cases, reached 100%, consistent with within-species relatedness (Fig. S3). When similar isolates (ANI values) were clustered hierarchically, two large clusters were discernible (Fig. S4). The larger cluster (in orange) comprised highly similar gut-derived lineages (connected by short branches), with human and animal gut samples intermingled (Fig. S4). Short branches among gut isolates point to a recent divergence, possibly from a common ancestor. We later revisit our ANI-based inference using phylogenetic analysis based on orthologous genes (See next section). Since gut isolates from animals were overrepresented due to a longer history of research (Fig. S4), we removed nearly identical isolates (ANI 99-100%) and used a rarefied subset for all downstream analyzes. In other words, *M. elsdenii* genomes were dereplicated at a ≥99% Average Nucleotide Identity (ANI) threshold, resulting in smaller number of representative isolates that capture the diversity of the larger number of genomes. ANI-based clustering analysis on the rarefied (dereplicated) subset (37 samples consisting of gut *M. elsdenii* and *Megasphaera* species from vagina) reproduced the tree structure identifiable in the full dataset (Fig. 1). To assist our downstream analyzes, we labeled three weakly differentiated gut-derived lineages as “gut_1”, “gut_2”, and “gut_3”. *Megasphaera* species associated with the human vagina dominated the second heterogeneous branch (in green), subdivided into three sub-branches: the first “mixed” group was composed of *M. massiliensis, M. vaginalis, M. butyrica, M. hexanoica, M. hominis*; the second group (“vag_1”) represented by vaginal phylotype 1, and the third group (“vag_2”) contained vaginal phylotype 2 as well as *M. hutchinsoni* (Fig. 1). As expected, *Megasphaera* species associated with the vaginal niche were more divergent and heterogeneous than gut lineages of *M. elsdenii*. The extent of dissimilarity between the two vaginal groups, “vag_1” and “vag_2”, was significantly higher (longer branches) than that observed between gut isolates from different hosts (human,– swine, and cattle); however, see the phylogenetic analyzes below. It is notable that *M. elsdenii* populations (lineages) from human fecal samples belong to a genetically homogeneous cluster, which is composed of isolates from ruminant (cattle) and non-ruminant (swine) animals (Fig. 1). Furthermore, gut lineages from different organisms show no clear indication of differentiation into host-specific clusters.

**FIG 1.**
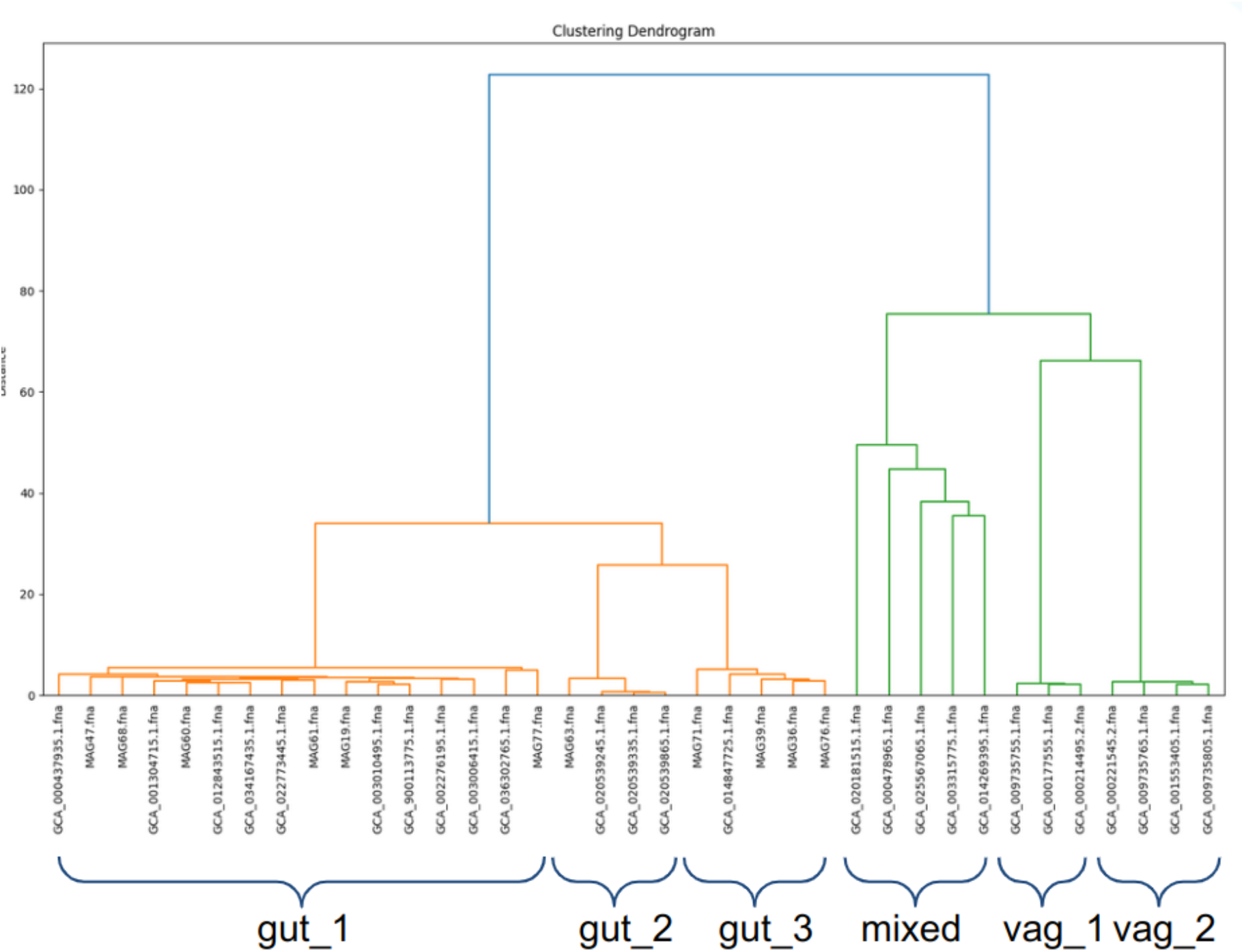
Hierarchical clustering tree of 37 (rarefied subset) samples based on average nucleotide identity (ANI).

### Phylogenetic analysis based on orthologous genes

To better understand phylogenetic relationships between *M. elsdenii* lineages and related species, we used single-copy orthologs and constructed a rooted phylogenetic tree. Our phylogenetic analysis included the rarefied subset of 37 *M. elsdenii* lineages and related *Megasphaera* species. We also included more distant taxa, such as *Dialister*, *Veillonella* (from the human gut and oral cavity), and *Negativicoccus* (from the human gut), as well as *Clostridium botulinum* as an outgroup (Fig. 2).

**FIG 2.**
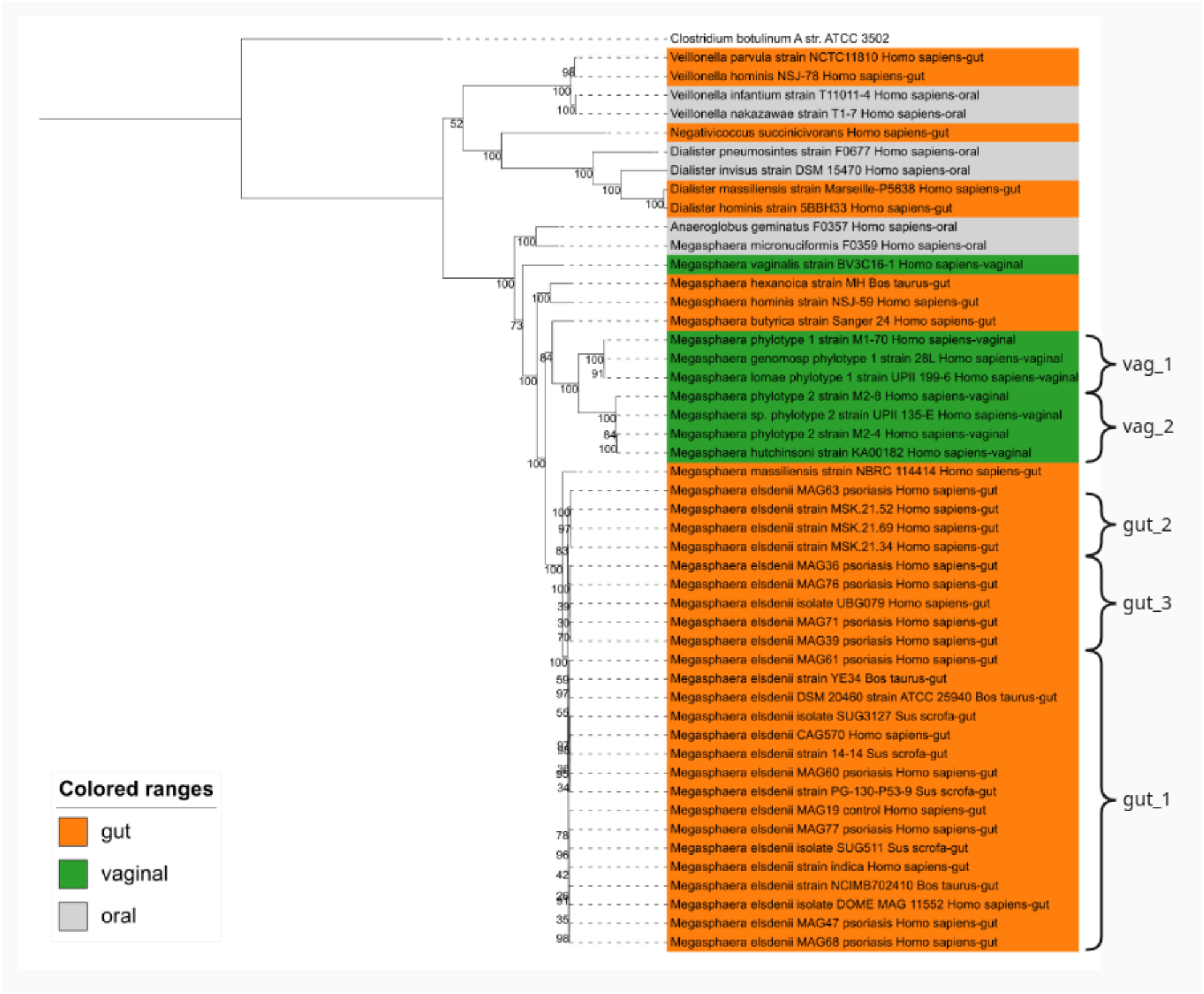
Phylogenetic tree of *M. elsdenii* and related taxa based on orthologous genes. *Megasphaera* samples from gut — orange, human vagina — green, oral cavity — gray; *Clostridium botulinum A str. ATCC 3502* represents an outgroup. A phylogenetic tree was constructed using the LG+I+G4 model.

Observed divergence of single-copy orthologs implies that *M. elsdenii* and other *Megasphaera* species are well separated from related genera, such as *Dialister* and *Veillonella*. Of note is a single sample of *Anaeroglobus geminatus* placed together with *Megasphaera micronuciformis* but well outside other branches, representing most *Megasphaera* species (Fig. 2). Thus, whole-genome analysis seems to support earlier inferences based on rRNA (See Figure 2 in [23]), which placed *Anaeroglobus geminatus* within the *Megasphaera* branch and suggested reclassification into *Megasphaera*. For *Megasphaera* species, our phylogenetic analysis recapitulated ANI-based inferences (See Fig. 1) on the major division between species associated with the human vagina and gut, and relatively weak divergence between *M. elsdenii* gut-associated lineages (Fig. 2).

Indeed, half of the human gut lineages are intermingled with the gut isolates from animals. Furthermore, the two arbitrarily defined groups of human gut lineages, designated as gut_2 and gut_3 (to assist with description), are only weakly diverged (very short branches) from animal gut isolates. Similarities summarized at the genome scale and reflected in phylogenetic trees can obscure smaller-scale changes at individual genes with important metabolic functions.

### Constructing inter-species pan genome for *M. elsdenii* gut isolates and vagina-associated *Megasphaera* species

We first constructed a pan-genome for *Megasphaera* species to broadly assess the proportion of genes that *M. elsdenii* gut lineages share with *Megasphaera* species that associate with different niches - the human vaginal niche. We grouped samples based on their phylogenetic relationships and visualized shared genes using a heatmap (Fig. 3A) [41]. Blue bars indicate shared genes, and white spaces indicate missing genes. It is obvious that *Megasphaera* species associated with the vaginal niche have a drastically different genetic makeup compared to the gut-associated *M. elsdenii* lineages. Most of the inter-species pan-genome is absent in vagina-associated species (Fig. 3A). In contrast, gut-associated lineages from different hosts show no noticeable differences in gene content at this crude scale of assessment. It is likely that the large differences observed between vaginal species and gut lineages obscure subtler differences that might exist between gut lineages from different hosts. We, therefore, removed *Megasphaera* species associated with the human vagina when we attempted to predicts delineate metabolic features between *M. elsdenii* gut lineages from different host.

**FIG 3.**
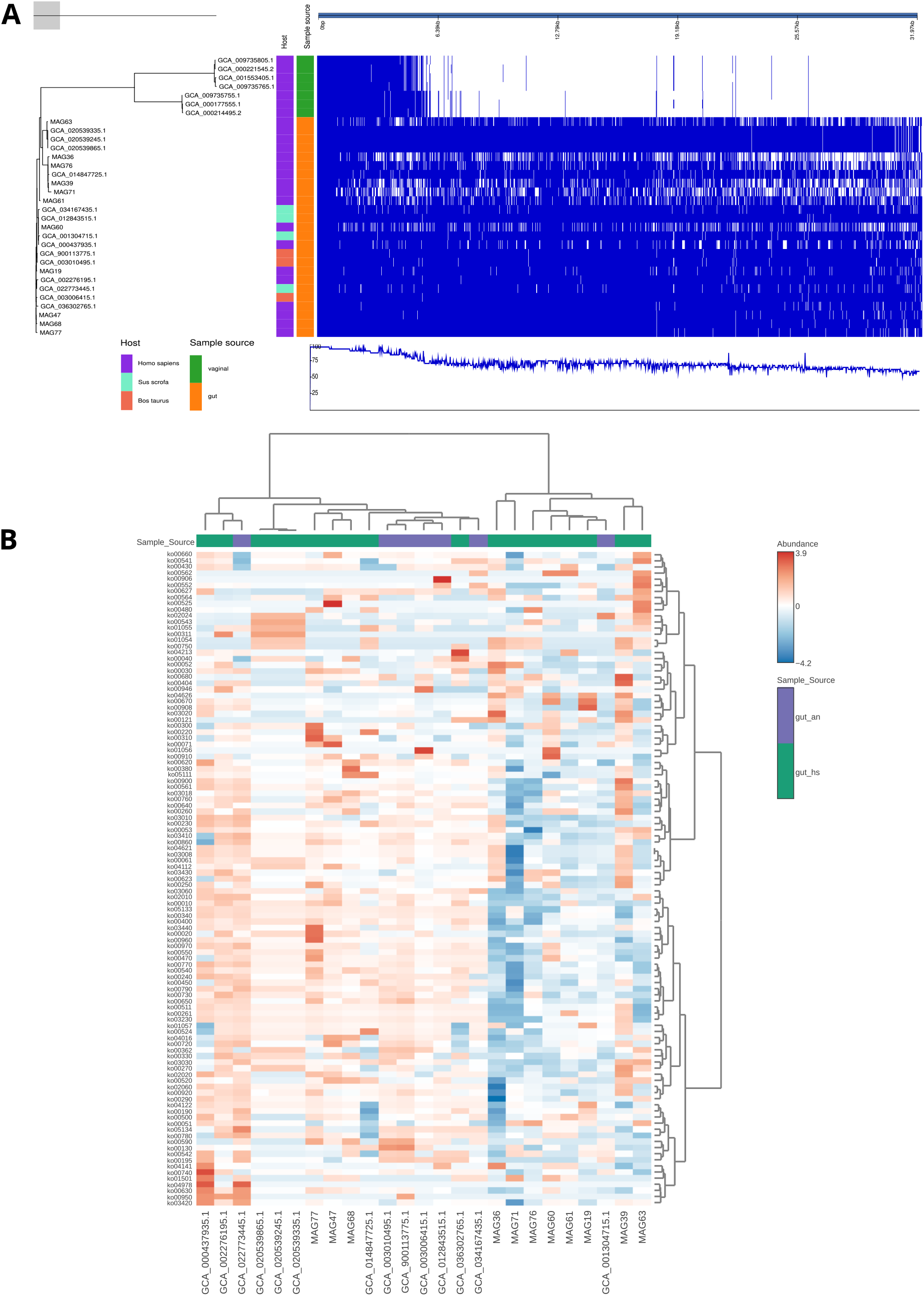
Distribution of genes and pathways among *M. elsdenii* gut lineages and *Megasphaera* species associated with the human vaginal niche. (A) Gene presence-absence analysis of the inter-species pan-genome. Heatmap visualization is based on the Phandango tool, which integrates Roary-based pangenome inference for *M. elsdenii*, metadata such as host origin and niche association. (B) Heatmap representing KEGG Pathway abundance with row and column clustering (Distance-based Ward clustering).KEGG pathway abundances were converted to distances (Jensen–Shannon divergences) and clustered using Ward’s method.

### Analysis of KEGG pathways and CAZymes in gut lineages of *M. elsdenii*

To translate the observed gene content into microbial functions, we mapped genes to KEGG metabolic pathways. We organized KEGG pathways into heatmap rows and examined how predicted metabolic profiles for *M. elsdenii* gut lineages corresponded to host associations (Fig. 3B). In the resulting heatmap, similarities in KEGG pathways were used to cluster *M. elsdenii* gut samples into metabolic profiles (the Ward clustering tree above the heatmap), and each sample was annotated according to host association: gut_hs - human gut and gut_an - animal gut. As shown, gut samples from humans and animals did not form clear-cut clusters of metabolic profiles driven by host association. To detect subtle trends in KEGG pathway variation, we also applied a more sensitive approach: principal component analysis. Neither PC1, capturing 23% of KEGG pathway variation, nor PC2 (13% of KEGG pathway variation) separated gut samples from humans and animals (Fig. 4A).

**FIG 4.**
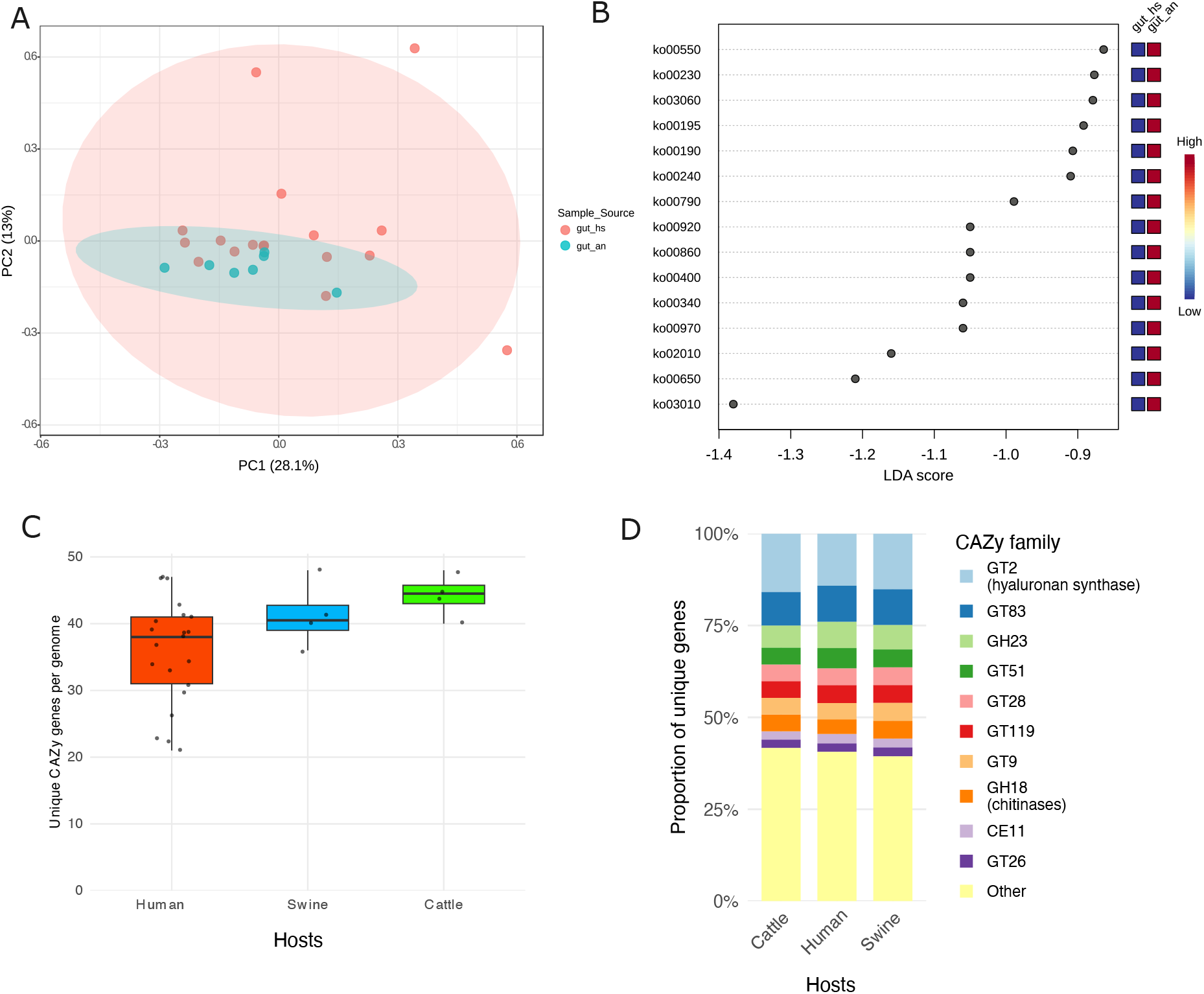
KEGG pathways and CAZymes in *M. elsdenii* gut lineages. (A) The top two principal components of KEGG pathway variation in the combined set of *M. elsdenii* genomes from human and animal gut. (B) KEGG pathways ranked by discriminatory power (LDA scores) to separate human and animal gut samples of *M. elsdenii*. Based on linear discriminant analysis (LEfSe). (C) Total number of genes from the CAZy database in *M. elsdenii* gut lineages grouped hosts. (D) The proportion of carbohydrate-active enzyme families in *M. elsdenii* gut lineages. Data shown for the top 10 most frequent families: GT - glycosyltransferases, CE - Carbohydrate esterases, GH - glycosyl hydrolases.

While heatmap representations, clustering, and PCA are visually appealing, the differences among gut lineages may be too subtle to drive unsupervised clustering (here, Ward clustering and PCA). We therefore turned to a supervised LEfSe approach [58] and searched for differences between predefined classes of interest — human gut and animal gut. LEfSe analysis ranked several pathways, but their small LDA-scores (less than 2) suggested relatively weak differential abundance between human gut and animal gut samples (Fig. 4B). In other words, if we consider individual pathways, each with a small effect size (LDA scores ranged from 0.9 to −1.4), they would not be informative for distinguishing gut lineages between humans and animals.

The *M. elsdenii* genomes analyzed in this study did not have systematic differences in KEGG pathways between hosts, despite their host organisms having different digestive tracts and diets. An earlier study showed differences in the gene repertoire for carbohydrate-degrading enzymes between a ruminal isolate and two human gut isolates [11]. Given this prior evidence, we asked whether gut lineages of *M. elsdenii* from humans, swine, and cattle have systematic differences in gene repertoire for degrading carbohydrates. Using the CAZy database as a reference, we searched for homologous genes encoding carbohydrate-binding and degrading enzymes. On average, *M. elsdenii* from humans, swine, and cattle had 36, 41, and 44 CAZyme-encoding genes (Fig. 4C), but differences were not statistically significant (p-values ranged from 0.12 to 0.59, pair-wise Wilcoxon tests). It is important to note that four genomes (dots in the boxplot, Fig. 4C) for swine and four for cattle represent diversity of dozens of nearly identical isolates (ANI 99-100%) that were dereplicated at a 99% ANI (See diversity of the full dataset in Fig. S3 and Fig. S4). Next, we classified genes into CAZy families, but their proportions did not differ between different hosts based on ANOVA (Analysis of Variance) with Tukey’s Post-Hoc Test (p-values ranged from 0.17 to 1) (Fig. 4D). Thus, our analyses suggest that gut-associated lineages of *M. elsdenii* when grouped by host (human, swine, or cattle) do not show consistent group-level differences in the number of CAZymes.

We explored the distribution of CAZy gene families in more detail to understand discrepancies with earlier findings based on three genomes [11]. A bubble plot was generated to visualize how individual genomes differ in the number of detected genes in CAZy families (Fig. 5). To spot discrepancies, the human gut isolate (GCA_002276195.1) from the earlier study by [11] was highlighted using the green horizontal rectangle and three ruminal isolates with orange horizontal rectangles. It should be noted that these three ruminal isolates are nearly identical (ANI 99-100%) to the type strain analyzed used in [11], which was present in our full 98 genome set. First, the number of encoded CAZyme genes varied across genomes within each host group (Fig. 5). These individual differences are not shared by all group members (when grouped by host) and therefore did not appear as systematic group-level differences in our statistical analyses. Next, we highlighted CAZy families (GH25, GH32, GH43, GH53, GH73, and GH77) previously reported by [11] as differentially present in human gut isolates but absent in the type strain (ruminal isolate). Half of these gene families were either absent (GH32, GH53, and GH73 were not detected) or showed no difference (GH25, GH43, and GH77) when detected. Finally, each genome was classified according to MIMAG standards as high-quality (HQ) or middle-quality draft, based on the quality of the genome assembly: completeness and contamination scores computed by checkM for MAGs (Table S4) and BUSCO scores for published genomes (Table S3). It can be seen that some middle-quality drafts (MAG36, MAG39, and MAG76) tend to have fewer annotated genes. On the other hand, the number of detected gene families also varies among high-quality drafts. In addition, some high-quality MAGs, such as MAG61, consistently have fewer genes in certain gene families (GH23, GT51, GT119, and GT28) than middle-quality MAGs, such as MAG39, MAG36, and MAG63. In sum, observed variation in the number of detected genes most likely reflects genetic variation within bacterial species, but some gene-missingness patterns may be due to genome quality.

**FIG 5.**
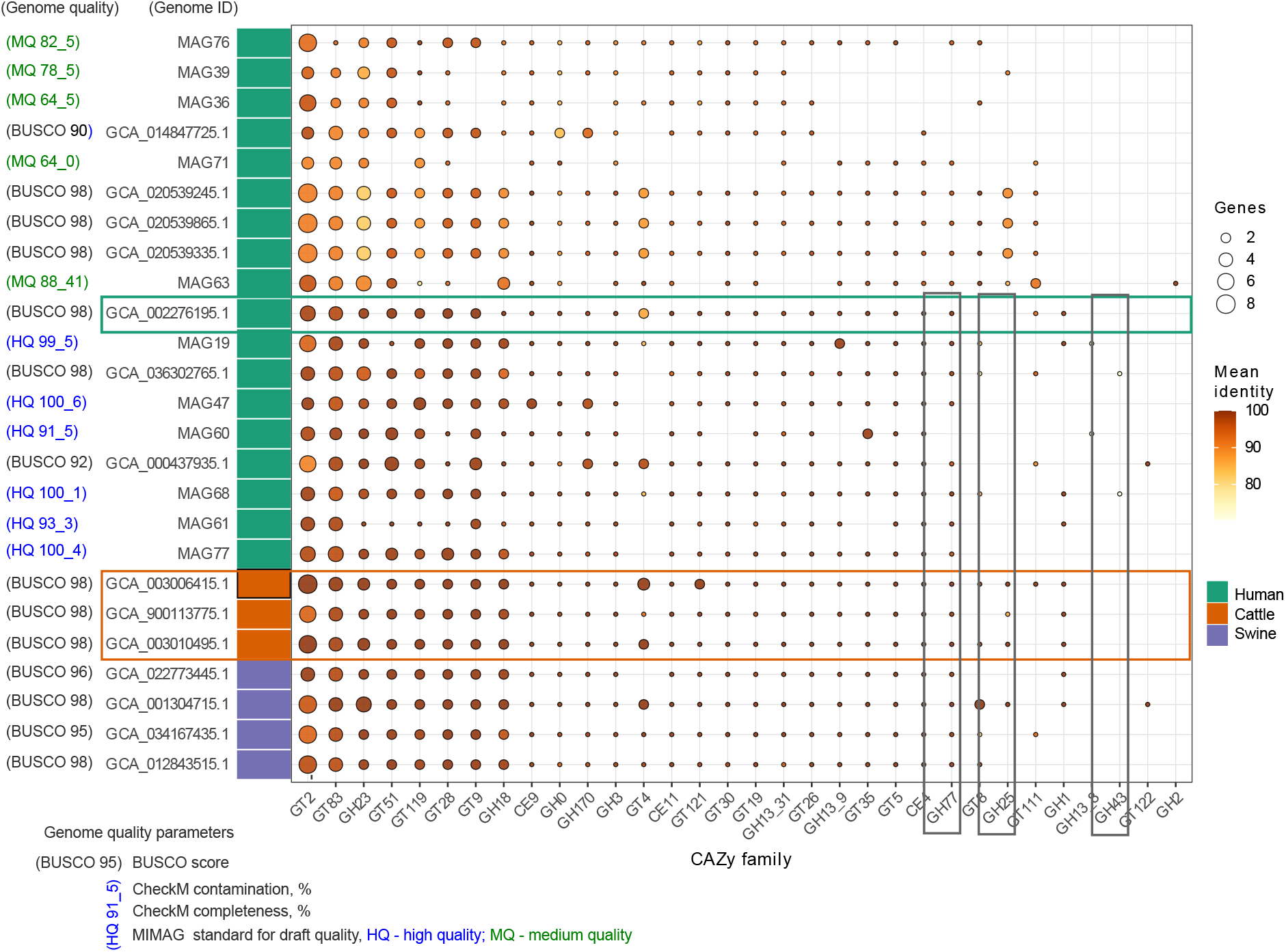
Number of CAZy gene families across *M. elsdenii* genomes. Abbreviated CAZy family names are listed along the x-axis. Circle size indicates the number of homologs from a particular family, and the color corresponds to the mean identity with the known reference sequence.

### Nutrient requirements predicted based on genome-scale metabolic model

In this section, we attempted to predict the nutrients required for *M. elsdenii* growth, focusing on carbon and energy sources. For each assembled genome, we reconstructed a genome-scale metabolic model using the gapseq tool [50]. The inferred metabolic models were used to predict which organic and inorganic compounds are required for bacterial growth under anaerobic conditions (Fig. 6). Organic compounds predicted by gapseq could be classified into three potential sources of carbon and energy: carbohydrates, organic acids, and amino acids. For example, gapseq predicted that all gut lineages of *M. elsdenii* require six amino acids (e.g., L-tyrosine, L-serine, L-phenylalanine, L-glutamine, and L-arginine) to grow. However, in most bacteria, amino acids are not the first-choice source of energy; they are used as building blocks for cellular components. According to gapseq predictions, some gut lineages from human and swine (highlighted by dashed rectangles) required a wider repertoire of amino acids and chorismate for growth (Fig. 6). By highlighting seven such gut lineages, we noted that four (green dashed rectangle) corresponded to MAGs with a middle-quality draft, two (blue dashed rectangle) to high-quality draft, and one (black dashed rectangle) to a high-quality published genome. We note that MAGs, classified as middle-quality, tend to be overrepresented (4 out of 5 middle-quality MAGs) in this set of highlighted genomes.

**FIG 6.**
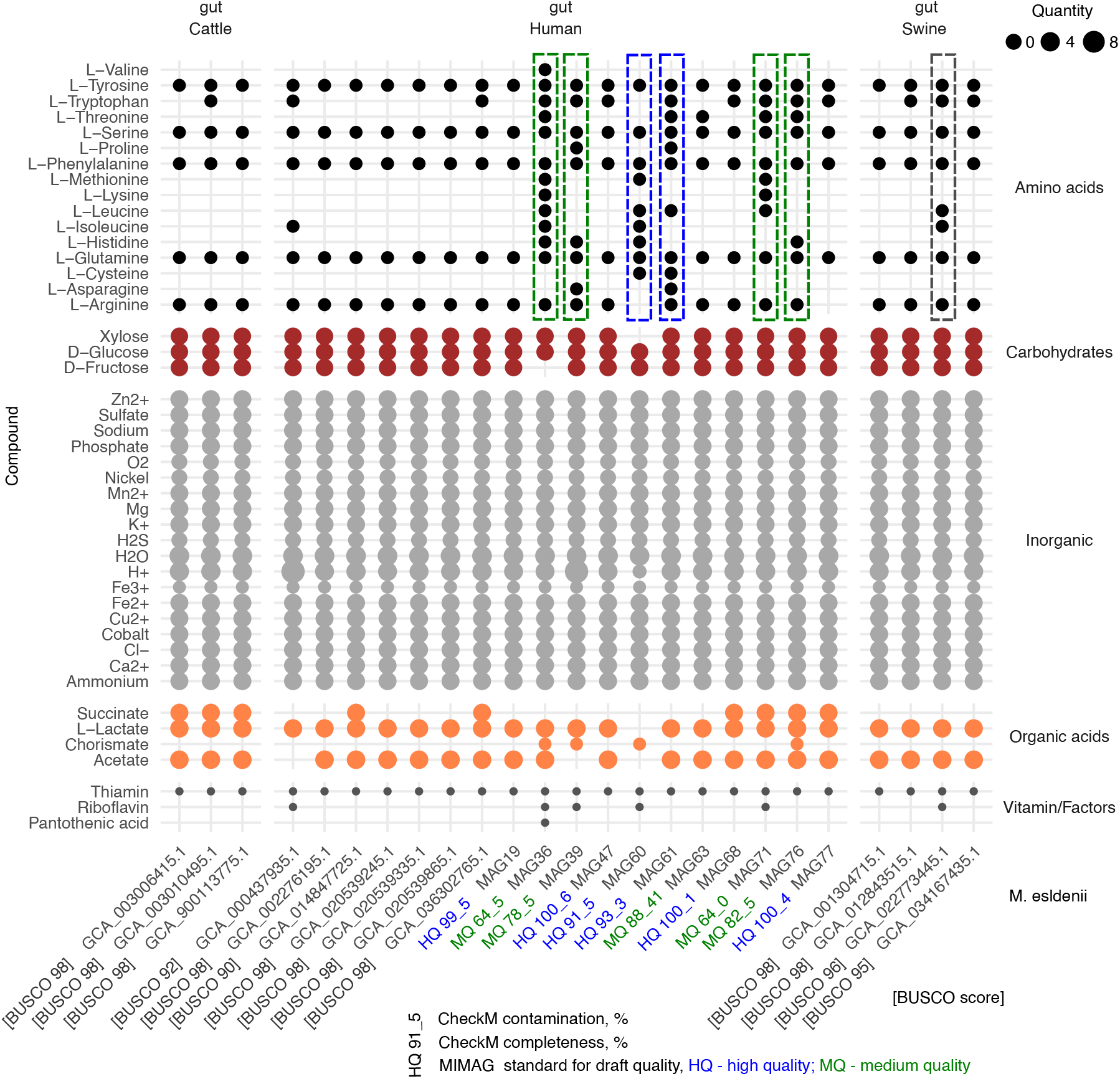
Growth media compounds predicted based on the genome-scale metabolic model. Based on gapseq analysis.

We next focused on carbohydrates and their fermentation products, as they represent the preferred energy source. Gapseq predicted that three simple carbohydrates (xylose, D-glucose, and D-fructose) can be consumed by *M. elsdenii* gut lineages. Importantly, all *M. elsdenii* gut lineages had the potential to utilize key fermentation products derived from carbohydrates, acetate, and L-lactate, regardless of host association (Fig. 6). Some gut lineages (9/25) had the potential to utilize succinate, and only a few (4/25) had the potential to utilize chorismate. Nevertheless, we were not confident in chorismate, as it was predicted for only three MAGs, all of which corresponded to middle-quality drafts, and one MAG was classified as a high-quality draft. In sum, it appears that gut lineages of *M. elsdenii* from different hosts can consume both simple sugars and secondary fermentation products, and none of them lost genes needed to utilize acetate and L-lactate. Finally, we tested whether human gut lineages possess the same set of genes involved in lactate uptake and utilization as those in better-studied animal gut isolates. Specifically, we screened for lactate permease (lutP), which imports lactate into the bacterial cell, and for genes in the conserved *lutABC* operon that encode the oxidative conversion of L-lactate to pyruvate. We found that all the analyzed gut samples, regardless of origin (human gut or acidic rumen), carry the lactate permease gene, and the lutC, lutB, and lutP genes of the lutABC operon.

### Lack of genes for known virulence factors

Even if *M. elsdenii* in the human gut grow in response to lactate accumulation, we wondered whether they encode genes associated with a detrimental impact on the host. For example, genes for host cell adherence and enzymatic degradation of host cell components. Using the MMseqs2 tool [48], we searched for genes that are homologs to known virulence factors in the VFDB 2022 database [51]. We did not find close homologs of known virulence genes using a sequence-similarity ≥ 70%. Because analyses based on close sequence similarity can miss structurally similar proteins, we also applied VirulentHunter, a pre-trained ESM2 protein language model [53] fine-tuned on virulence genes [54]. We refer to VirulentHunter findings as predicted virulence factors (pVFs), assuming that they represent structurally similar proteins to virulence factors. We ran VirulentHunter by including one known gut commensal, *Dialister hominis*, and one known pathogen, *Clostridium botulinum*, which are phylogenetically related to *M. elsdenii*. We found that *M. elsdenii* samples from human gut (divided into healthy donors and psoriasis patients) had a comparable number of pVFs as a known gut commensal, *Dialister hominis* (Fig. 7A). Next, both *M. elsdenii* and *D. hominis* had a significantly lower (ANOVA with post-hoc Tukey test, p < 0.05) number of pVFs compared to a true pathogen, *Clostridium botulinum*. Moreover, *Clostridium botulinum* had statistically higher numbers of pFVs from multiple functional classes, such as adherence, effector delivery system, exotoxin, immune modulation, and motility (Fig. 7B). In sum, the protein language model predicted a larger number of pVFs for a true pathogen than for known commensal and *M. elsdenii*, but predictions are likely noisy and useful only as an exploratory step.

**FIG 7.**
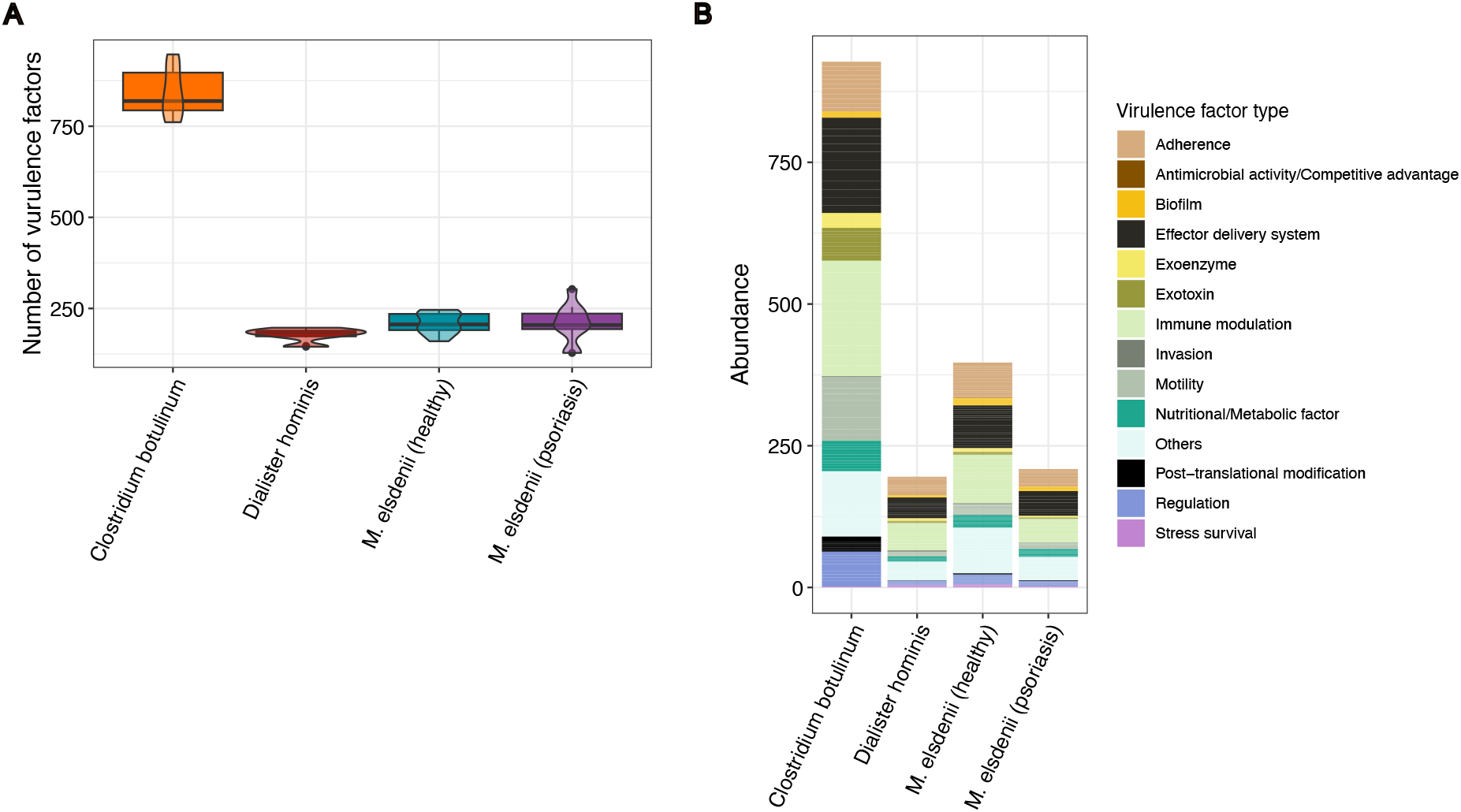
The distribution of virulence determinants predicted with VirulentHunter in *Megasphaera* genomes isolated from healthy donors and psoriatic patients, coupled with positive (*Clostridium botulinum*) and negative controls (*Dialister hominis*). (A). The mean genome-wise amount of virulence factors found in the bacterial groups. (B) Total proportion of functional groups of virulence determinants classified according to the VirulentHunter utility.

## DISCUSSION

Gut bacterial species are suspected of being detrimental if increased in abundance and correlate with inflammatory conditions, but such correlations can have alternative explanations. For example, patients can develop an altered gut environment due to low-grade inflammation, and commensals specialized in inflammation by-products can benefit from such changes [59]. In this study, we focused on *Megasphaera elsdenii*, which has repeatedly been shown to be disease-associated in many human inflammatory conditions. We were motivated by the recent evidence that *M. elsdenii* relative abundance linearly correlates with increasing fecal calprotectin in both patients and healthy controls [17], and that animal gut isolates also increase in relative abundance in dysbiotic rumen to consume lactate [60, 61]. Indeed, fecal calprotectin indicates immune cell effector activity in the intestine [62], and by-products of inflammation, such as lactate, can provide fitness advantages for certain bacteria that benefit from lactate and other inflammation by-products [63]. In light of this alternative view, our first objective was to understand genetic variability and relatedness among *M. elsdenii* lineages from the human gut and the animal rumen.

### Genetic proximity of human and animal gut isolates

Previous phylogenetic analyzes based on whole genomes placed two human-derived isolates on a separate branch next to the type strain from cattle rumen [11]. It is unlikely that three isolates from two organisms can adequately cover the genetic diversity of the bacterial species. In this study, using a larger sample of genomes (a full dataset of N=89 and a dereplicated subset of N=37), we show that human gut populations of *M. elsdenii* do not form a separate branch. Instead, they all belong to the same branch as animal isolates, which, in addition, represent a highly homogeneous clade (Fig. 1). Moreover, we expected that human gut-associated lineages would be differentiated from ruminal isolates and, at the same time, fall closer to swine isolates due to similarities in the digestive tracts [64]. Instead, we found that human gut lineages intermingle with cattle and swine isolates, and there is no clustering according to host association (Fig. 2). Based on the high genomic identity between *M. elsdenii* lineages from humans and animals (as evidenced by high ANI values and clustering patterns), we hypothesized that they have a recent common origin. Our genome-based inference agrees with the earliest phylogenetic analyzes based on rRNA gene variation. Namely, earlier rRNA gene-based study found that gut-derived isolates were highly homogeneous and proposed that *M. elsdenii* strains most likely expanded recently from a common ancestor [65]. Thus, our phylogenetic analysis (Fig. 2) and ANI-based clustering support the idea that *M. elsdenii* from human and animal gut likely derive from a recent common ancestor, as proposed earlier [65].

### *Megasphaera* vaginal phylotypes lack most of the *M. elsdenii* pan-genome elements

We added *Megasphaera* species from the human vagina to our analyses to represent a related species from a distinct niche than the animal intestine. We found that *Megasphaera* species from the same vaginal niche accumulated more genetic divergence (longer branches on phylogenetic trees) than *Megasphaera elsdenii* lineages derived from different host organisms with distinct diets and gastrointestinal tracts: humans, swine, and cattle. Inter-species pan-genome analysis showed that vagina-associated species lack many of the genes present in *M. elsdenii* gut lineages (Fig. 3A). It was apparent that vagina-associated species would show strong differentiation from gut lineages in terms of encoded metabolic pathways, and that this dominant signal would overshadow any subtle differences between gut lineages of *M. elsdenii*. To focus on subtle differences between *M. elsdenii* gut lineages in terms of metabolic profiles, we decided to exclude vagina-associated species and performed metabolic predictions by focusing on *M. elsdenii* gut lineages.

### *M. elsdenii* gut lineages from different hosts lack host-specific enrichment of carbohydrate-degrading enzymes

A previous comparative genomic study used two human fecal isolates and showed that they have a broader spectrum of carbohydrate-active enzymes than a ruminal isolate, the type strain *Megasphaera elsdenii* (DSM20460) [66]. These findings looked reasonable given that cattle microbiota rely on plant carbohydrates [67]. In contrast, humans like swine are omnivorous, and therefore their gut microbiota is expected to encode a wider repertoire of carbohydrate-degrading enzymes [68, 69]. Nevertheless, based on a more representative sample of *M. elsdenii* from different hosts, we found no evidence (Fig. 4 and Fig. 5) for a wider repertoire of CAZy enzymes among human gut lineages compared to ruminal isolates. We hypothesized that prior evidence most likely reflected individual differences between three genomes available at the time [11], i.e., two human gut isolates and one type strain from cattle [11]. By including previously published human gut isolate (GCA_002276195.1) from [11] and ruminal isolates that were identical (99%-100% ANI) to the type strain *Megasphaera elsdenii* (DSM20460), we explored how *M. elsdenii* genomes varied in terms of detected CAZy genes (Fig. 5). We found that the number of detected CAZy genes (binned by gene families) among *M. elsdenii* gut lineages can vary within the same host and between host groups (Fig. 5). However, observed gene presence/absence patterns were not shared among all members of the host-associated groups (humans, swine, or cattle) and therefore did not manifest as a group-level feature in our statistical tests (Fig. 4). While differences in gene content between genomes can sometimes stem from incomplete assembly, observed variability likely reflects true within-species diversity, typical of microbial populations. In sum, earlier findings by [11] are likely to reflect features of individual isolates (Fig. 5) and are not generalizable to larger samples from the human gut and the cattle rumen (Fig. 4).

### Nutrient requirements of *M. elsdenii* predicted from genome-scale metabolic modes

In the current study, we used genome-scale metabolic models to predict the nutrients required for microbial cell growth. Specifically, we performed metabolic predictions for 21 *M. elsdenii* lineages from human gut, four from cattle, and four from swine using the gapseq tool. It is reassuring that predicted nutrient requirements, such as potential to synthesize amino acids and vitamins, were consistent with previous study findings on human gut isolates [66]. For example, Shetty et al. [66] showed that *Megasphaera elsdenii* genomes from their human gut isolates had genes for the synthesis of essential amino acids, such as histidine, lysine, methionine, threonine, and tryptophan. Our predictions using gapseq suggested that human and animal gut *M. elsdenii*, with few exceptions, also have the potential to synthesize these amino acids and therefore did not require histidine, lysine, methionine, or threonine in the predicted growth medium (Fig. 6). Next, we found that *M. elsdenii* from the human and animal gut encode genes for pathways (and potential) to utilize three simple sugars (xylose, D-glucose, and D-fructose) and two organic acids, L-lactate and acetate (Fig. 6). Based on the predicted nutrient requirements, human gut *M. elsdenii* lineages, like animal gut isolates, can be placed among secondary/intermediate consumers in the microbial cross-feeding chain in the anaerobic gut. More precisely, these gut lineages likely perform primary fermentation (of simple sugars) and secondary fermentation (of lactate and acetate). This metabolic specialization likely explains why they do not have differences in CAZy gene families depending on the host diet (cattle versus humans), which would place them upstream in the cross-feeding chain among gut bacteria, such as Bacteroides spp. and Prevotella spp., that degrade complex carbohydrates [70]. Next, our predictions about lactate utilization for human gut lineages are consistent with recent in vitro experiments with *M. elsdenii* derived from human feces. Namely, microbial community from feces produced more valerate (and propionate or butyrate, depending on the conditions) from lactate when *M. elsdenii* was present in the batch culture [12]. In addition, *M. elsdenii* strongly co-occurred with the lactate-producing genus Lactobacillus.

While metabolic predictions in our study suggest that *M. elsdenii* from the human gut can utilize lactate, genomic data, and inference methods used can not distinguish catabolite repression mechanisms and inform on the preferences between organic acids (lactate and acetate) and simple sugars. So far, lactate preference over glucose has been experimentally shown only for ruminal isolates [23]. Finally, as far as we can tell from metabolic predictions, succinate, a central metabolic intermediate, is dispensable for most studied gut isolates, except for ruminal isolates and some human gut lineages. In contrast, predictions about another metabolic intermediate, chorismate, may be a potential example of missingness due to incomplete genome assembly (Fig. 6). Only four gut lineages required chorismate, and three of them were MAGs, classified as middle-quality draft. On the other hand, it is reassuring that random missingness due to genome assembly, which is inevitable, did not substantially disrupt metabolic predictions, as genes for metabolic pathways essential for cell growth were present in all the studied gut lineages. Indeed, all the studied gut lineages from different hosts had pathways for the consumption of acetate and lactate (as well as the three simple sugars: xylose, D-glucose, and D-fructose) (Fig. 6). In sum, for the analyzed *M. elsdenii* gut lineages, lactate, as well as xylose, D-glucose, and D-fructose, seem to be conserved sources of energy, while acetate is known to be carbon source for this species.

### Is there evidence for detrimental function?

While *M. elsdenii* is considered a gut commensal and can grow in response to lactate; its relationship to the host’s health is poorly understood. Our analyses suggested that *M. elsdenii* from the human gut did not harbor close homologs to known virulence factors. This conclusion was based on the MMSeq2 search for close homologs (≥ 70% similarity) with known virulence factors. However, proteins can retain similar structure and function even after losing sequence similarity [71]. We therefore attempted to identify virulence factors based on structural similarity to known virulence factors. For that, we used a pre-trained ESM-2 protein language model in VirulenceHunter that was fine-tuned on virulence factors from three databases: VFDB 2022 [51], Victors [55], and PATRIC [56]. VirulenceHunter predicted that human gut *M. elsdenii* harbored ∼200 predicted virulence factors (pVFs), i.e., proteins structurally similar to known virulence factors. There is currently little empirical data to make sense of these kinds of pVFs. We note that the number of pVFs in *M. elsdenii* was comparable to that in a typical human gut commensal, *Dialister hominis*, but significantly lower (∼750) than for a true pathogen, *Clostridium botulinum*. To understand these findings, we considered the nature of the machine-learning approach and some context about the virulence factors. First, VirulenceHunter uses the ESM-2 model [72] that learns patterns governing protein structure and function [73]. Second, virulence factors evolve from proteins that serve non-pathogenic functions in free-living bacteria [74]. It is also known that not every bacterial strain carrying virulence factors can manifest pathogenicity unless it lands in a suitable tissue (niche) and in a host that is susceptible to that pathogen [75]. Hence, by taking into account that we did not find close homologs to virulence factors, we conclude that *M. elsdenii* contain structurally similar proteins to known virulence with numbers likely typical for commensals, not pathogens.

In sum, we find evidence that functional potential encoded in *M. elsdenii* genomes sampled from the human gut, including those of psoriasis patients, are similar to those in ruminal isolates. *M. elsdenii* from the human and ruminant animal gut likely fill the same trophic niche in the microbial cross-feeding chain in the anaerobic gut. Given the predicted metabolic profile, increased intestinal lactate in the inflamed patient’s gut may provide a plausible explanation for this species’ association with different inflammatory conditions. Below, we summarize what is known about gut inflammation in psoriasis patients to support our working hypothesis. We note that the eleven *M. elsdenii* samples assembled in this study were picked based on their strong correlation with fecal calprotectin concentration in psoriasis patients’ feces. Fecal calprotectin is a biomarker of intestinal inflammation [62], and it is known that intestinal inflammatory conditions are accompanied by increased lactate [76]. Psoriasis patients have evidence of low-grade inflammation [77], irritable bowel syndrome, and non-ulcerative colitis based on endoscopic examination [78], and it is known that experimentally induced psoriasis-like inflammation in mice also leads to intestinal inflammation [79, 80]. Therefore, in light of the existing evidence on the altered gut environment in psoriasis patients, we hypothesize that the correlation of this gut microbe with psoriasis is likely explained by *M. elsdenii* metabolic profile to consume lactate, along with acetate and simple sugars. So far, *M. elsdenii* gut isolates from humans showed no evidence for immunogenicity and have long been considered a gut commensal [10, 66, 12]. Possible side effects to the host, for example, connected with metabolic output, can not be excluded, but poorly studied. So far, experimental studies that reported detrimental microbial products haven’t traced them to *M. elsdenii* [81]. In conclusion, the gut commensal role for this species remains a more likely scenario, and there is currently no evidence that *M. elsdenii* contributes to psoriasis pathogenesis mechanistically.

## ACKNOWLEDGMENTS

This research was supported by the Russian Science Foundation grant number 25-15-00133 (Personnel Costs, Laboratory Supplies, Operational Costs). The authors, BY, ASh, KA, and VV acknowledge support from Saint-Petersburg State University research project 124032000041-1 (Computing, Transportation and Subsistence, Submission and Page Fees). RA was supported by the state assignment of the Ministry of Education and Science of the Russian Federation for IHNA & NPh RAS. The funders had no role in study design, data collection, interpretation, or the decision to submit the work for publication.

## DATA AVAILABILITY STATEMENT

The shotgun metagenomic data used in this study are available in the NCBI database under BioProject accession code PRJNA1102742. Previously published assembled genomes for *Megasphaera elsdenii*, *Megasphaera* vaginal phylotypes, *Clostridium botulinum*, and *Dialister hominis* are available in the NCBI database using GenBank Assembly IDs listed in Table S3.

## FUNDING

Russian Science Foundation, 25-15-00133, Milyausha Yunusbaeva, Bayazit Yunusbayev, Dina Sabirova

Saint Petersburg State University, project 124032000041-1, Bayazit Yunusbayev, Anton Shikov, Kirill Antonets, Veronika Vinichenko

Ministry of Education and Science of the Russian Federation, state assignment for IHNA & NPh RAS, Radick Altinbaev

## Supplemental material

Supplemental tables. Tables S1 to S4.

Supplemental figures. Fig. S1 to S4.

