## Supplemental Figures S1-S4 for "Genomic insights into the human gut commensal *Megasphaera elsdenii*: Relatedness and metabolic potential compared to animal isolates"

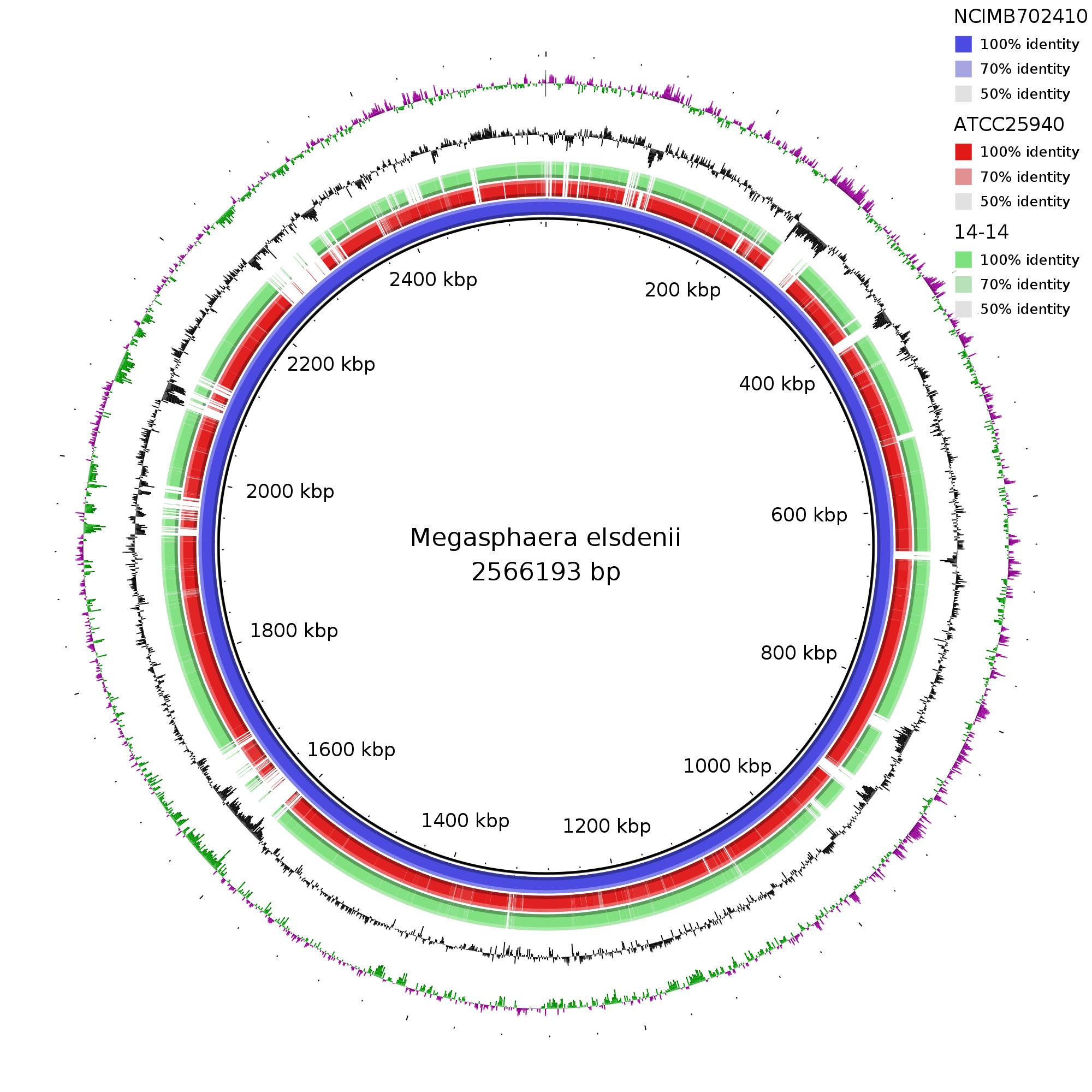
Fig. S1. *Megasphaera elsdenii* genome alignment using BRIG

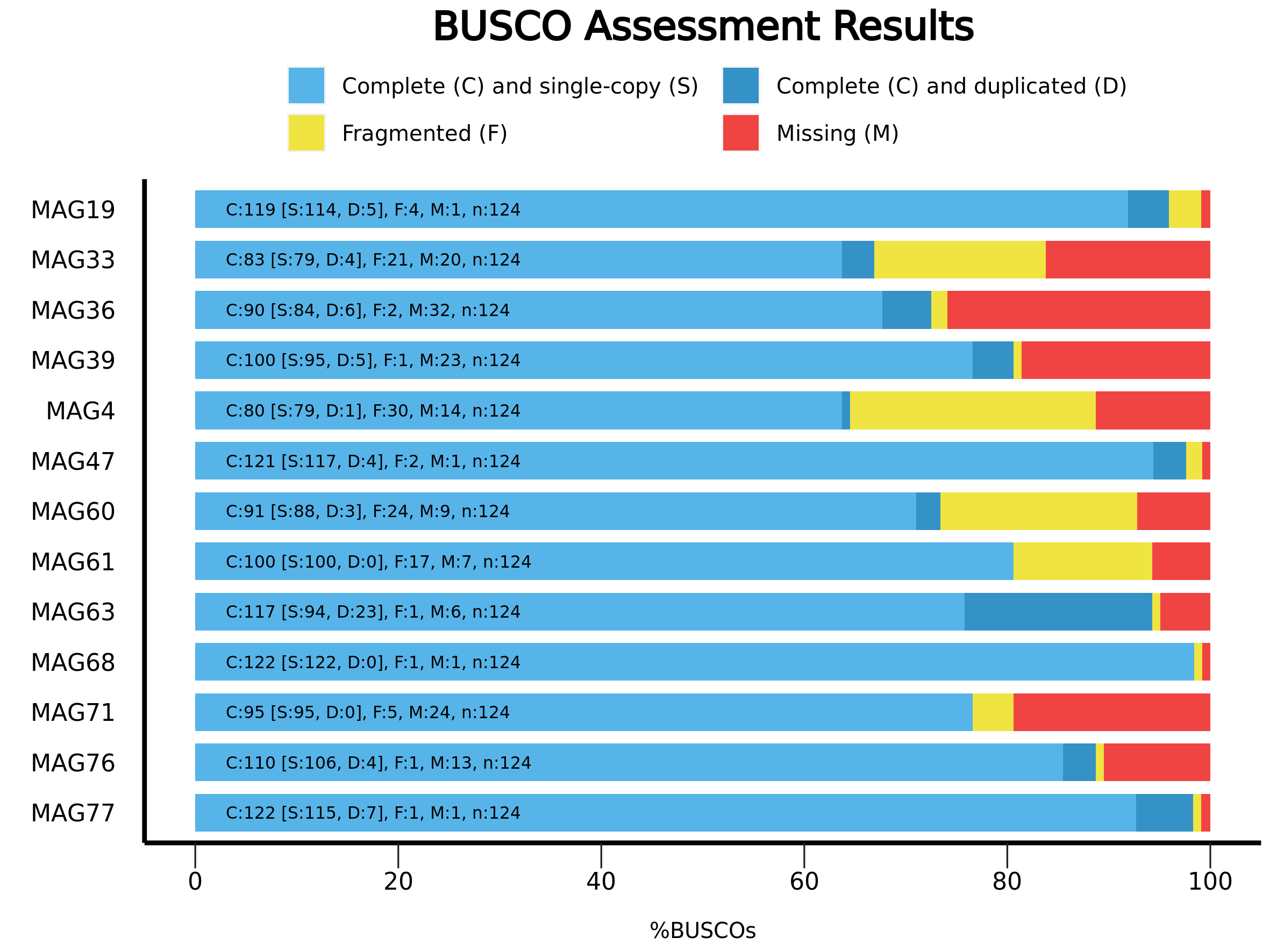

Fig. S2. Analysis of genome completeness for 13 MAGs using the BUSCO tool

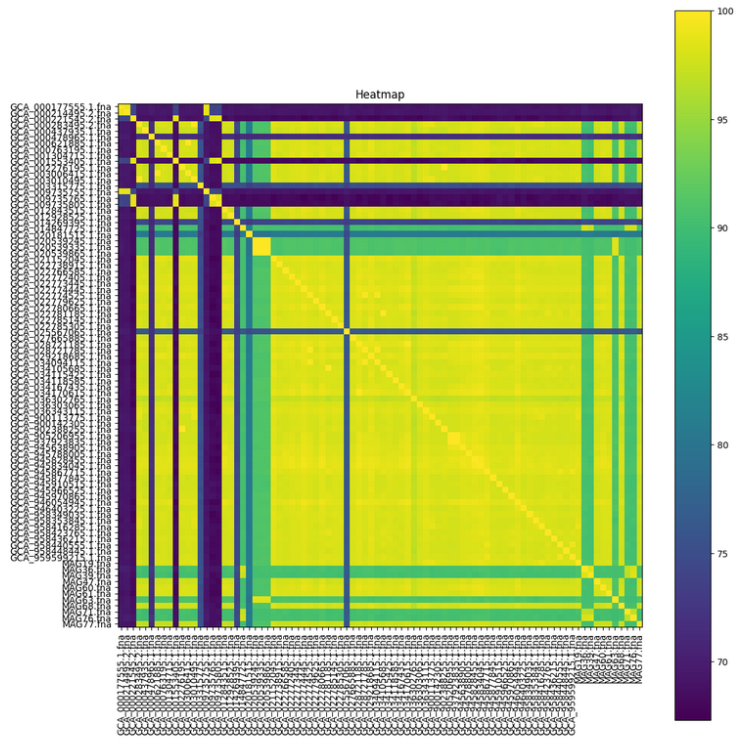
Fig. S3. ANI heatmap between 86 *Megasphaera* taxa

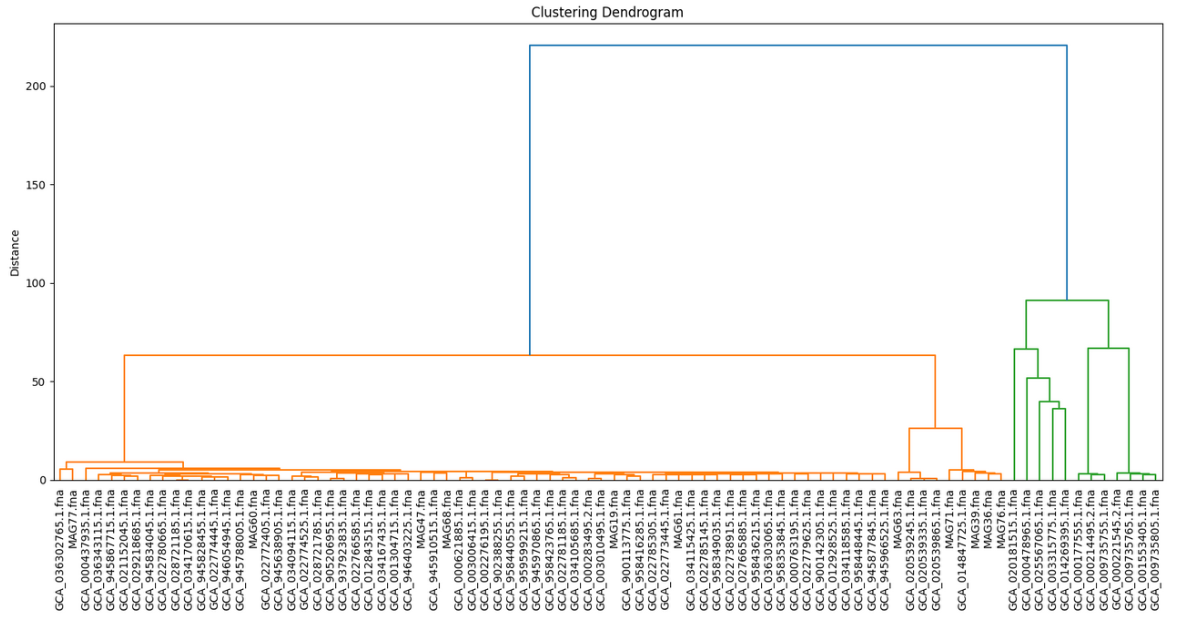

Fig. S4. Hierarchical clustering tree of 86 (full dataset) *Megasphaera* samples based on average nucleotide identity (ANI).
